# Non-invasive monitoring of drug action: a new live *in vitro* assay design for Chagas’ disease drug discovery

**DOI:** 10.1101/2020.01.15.907220

**Authors:** Anna F. Fesser, Olivier Braissant, Francisco Olmo, John M. Kelly, Pascal Mäser, Marcel Kaiser

## Abstract

New assay designs are needed to improve the predictive value of the *Trypanosoma cruzi in vitro* tests used as part of the Chagas’ disease drug development pipeline. Here, we employed a green fluorescent protein (eGFP)-expressing parasite line and live high-content imaging to monitor the growth of *T. cruzi* amastigotes in mouse embryonic fibroblasts. A novel assay design allowed us to follow parasite numbers over 6 days, in four-hour intervals, while occupying the microscope for only 24 hours per biological replicate. Dose-response curves were calculated for each time point after addition of test compounds, revealing how EC50 values first decreased over the time of drug exposure, and then leveled off. However, we observed that parasite numbers could vary, even in the untreated controls, and at different sites in the same well, which caused variability in the EC50 values. To overcome this, we established that fold change in parasite number per hour is a more robust and informative measure of drug activity. This was calculated based on an exponential growth model for every biological sample. The net fold change per hour is the result of parasite replication, differentiation, and death. The calculation of this fold change enabled us to determine the tipping point of drug action, i.e. the point immediately before the fold change becomes negative, independent of the drug concentration and exposure time. This time-to-kill over drug concentration revealed specific pharmacodynamic profiles of the benchmark drugs benznidazole and posaconazole.

**Author Summary:** Chagas’ disease, caused by *Trypanosoma cruzi*, is a chronic debilitating infection occurring mostly in Latin America. There is an urgent need for new, well tolerated drugs. However, the latest therapeutic candidates have yielded disappointing outcomes in clinical trials, despite promising preclinical results. This demands new and more predictive *in vitro* assays. To address this, we have developed an assay design that enables the growth of *T. cruzi* intracellular forms to be monitored in real time, under drug pressure, for 6 days post-infection. This allowed us to establish the tipping point of drug action, when the parasites stop multiplying and start to die. The resulting pharmacodynamics profiles can provide robust and informative details on anti-chagasic candidates, as demonstrated for the benchmark drugs benznidazole and posaconazole.

## Introduction

About 8 million people globally are infected with *Trypanosoma cruzi*, the causative agent of Chagas’ disease [1]. The progression of Chagas’ disease is divided into three phases: an acute, a chronic indeterminate and, in about 30% of the infected people, a symptomatic chronic phase. This last phase can begin decades after infection and is marked by severe cardiac or digestive symptoms. There are only two registered drugs available for Chagas’ disease, benznidazole and nifurtimox, and these suffer from severe side effects and variable efficacy [2]. Therefore, new treatment options are needed urgently.

Azoles like posaconzale and E1224, a prodrug of ravuconazole, were the most advanced drug candidates. However, in clinical trial, 80% of the posaconazole-treated patients relapsed within the 20 month follow-up after treatment, in contrast to 6% of the benznidazole-treated patients [2]. As a result, the research community was forced to rethink the pre-clinical drug discovery pipeline for Chagas’ disease [3, 4]. In particular, the design of *T. cruzi in vitro* assays had to be revisited to render them more predictive for the situation *in vivo*. A number of parameters were proposed for optimization: the choice of strains [5, 6], the life-cycle stages [7], the treatment regimens [8], and assay designs that assessed drug cidality. Tremendous progress has been made, especially in the area of high-content imaging technology for phenotypic assays [6, 8-14]. New wash-out designs were also introduced to assess reversibility and cidality of drug action [15, 16]. In combination with the development of more sensitive animal models [17-19], this permitted a focus on pharmacokinetic (PK) and pharmacodynamic (PD) parameters. PK-PD modeling allows treatment regimens to be modified for optimal exposure of the target organism to the drug candidate [20-22]. It also helps to define benchmark PK-PD parameters of the target product profile for drug candidates.

While there has been progress in modeling drug candidate PK profiles [23], the PD profile of a drug remains more difficult to determine. Two major aspects of PD need to be considered: time-to-kill and the question of whether drug action is concentration-driven or time-driven. Isothermal microcalorimetry has been used to determine time-to-kill for various pathogens, including African trypanosomes and *Plasmodium falciparum* [24, 25]. However, isothermal microcalorimetry cannot be used for intracellular amastigotes, the disease-relevant stage of *T. cruzi*, as it is impossible to differentiate the heatflow of the parasite from that of the host cell. Time-to-kill can also be determined by setting more than one temporal endpoint in an assay, which can mean one plate per time point has to be assessed [9, 16]. An efficient method to determine pharmacodynamic parameters still needs to be found.

Here, we describe a new *in vitro* live-imaging assay design and novel analysis methodology, which enables time-to-kill to be determined and identifies whether drug action is time- or concentration-driven.

## Methods

### Parasite and Cell cultivation

Mouse embryonic fibroblasts (MEF) were cultivated in RPMI supplemented with 10% FCS at 37°C, 5% CO_2_ and >95% humidity. MEF were sub-cultured once per week at a ratio of 1:10 after 5 min treatment with trypsin. *T. cruzi* STIB980 clone 1 (DTU TcI) was obtained from A. Osuna in 1983. Epimastigotes were maintained at 27°C in liver infusion tryptose (LIT) medium [26] supplemented with 20 µg/ml hemin and 10% FCS. Cultures were diluted weekly to maintain exponential growth. To stimulate metacyclogenesis, epimastiogte cultures were kept in the same medium for 3 - 4 weeks. About 10^7^ parasites from a predominantly metacyclic culture were taken to infect MEF for 48 h, and the cycle of amastigotes and trypomastigotes was maintained by infecting MEF weekly with an MOI of 1:1. eGFP-expressing parasites were kept for maximum of 4 weeks in the mammalian cycle.

### Transfection

Exponentially growing *T. cruzi* epimastigotes were synchronized for 24 h with 20 mM of hydroxyurea (Sigma) [27]. Following hydroxyurea removal by washing twice with PBS, 10^7^ epimastigotes were electroporated with 2.5 µg of pTRIX2-GFP plasmid (Fig 1 A) linearized with *Asc*I and *Sac*I (NEB). The plasmid had been derived from pTRIX-REh9 [28]. We used the Amaxa Nucleofector (programme X-014) with buffer Tb-BSF (Pacheco-Lugo et al., 2017), conditions we had found to be optimal for *T. cruzi*. 24 h after transfection, the parasites were diluted 1:10 in medium containing 100 µg/ml G418 (invivogen). Epimastigotes were cloned by limiting dilution. Clones were selected according to their eGFP expression level, infectivity and growth profile. Transgenic epimastigote cultures were maintained in the presence of 500 µg/ml G418.

**Fig 1.**
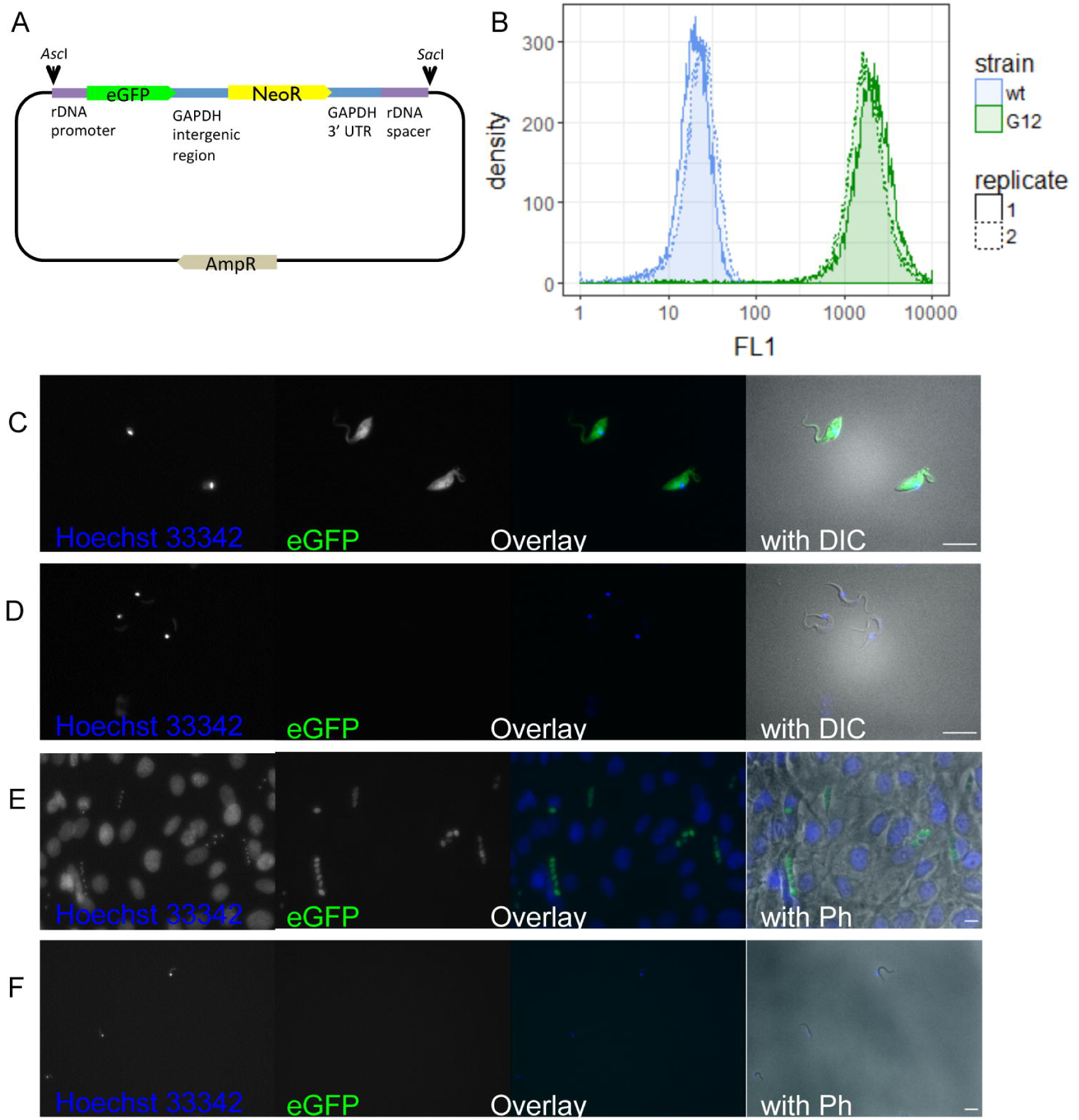
Transgenic parasites transfected with pTRIX2-GFP. (A) The plasmid pTRIX2-GFP was constructed by inserting the enhanced green fluorescent protein gene into the *T. cruzi* rDNA targeting plasmid pTRIX2-RE9h {Lewis, 2014 #70}. (B) Fluorescence levels of transfectant (G12) and wildtype (wt) epimastigotes were measured with flow cytometry. Epifluorescent images of transfectants in epimastigotes (C), metacyclic trypomastigotes (D), amastigotes in MEF (E), and trypomastigotes (F). Epimastigotes and metacyclic trypomastigotes were imaged with 630x magnification for 200 ms on DIC, 100 ms with the A4 filter cube (Hoechst 33342), and 300 ms with the L5 filter cube (GFP). Amastigote-infected Mef and trypomastigotes were imaged with 400x magnification for 50 ms with phase-contrast, 50 ms with the A4 filter cube (Hoechst 33342), and 815 ms with the L5 filter cube (GFP). Scale bars represent 10 µm.

### Flow cytometry

10^5^ epimastigotes in a small volume (approx. 100 µl) were fixed by the addition of the same volume of 10% formalin for 15 min at room temperature. After fixation, the volume was adjusted to 1.5 ml with PBS. Then, the parasites were analyzed for the levels of green fluorescence (FL1; excitation 488/10 nm and emission 530/30 nm) on a BD FACSCalibur (Becton Dickinson and Company) gating approximately from 100 to 2000 on the FSC and the SSC channel.

### Fluorescence microscopy

Epimastigote or trypomastigote parasites were deposited on a glass slide (Menzel Superfrost Plus), allowed to settle for 15 min, then washed with PBS and fixed with 10% formalin. For amastigotes, 10^4^ MEF per well were seeded on 16-well LabTek chamber glass slides (LabTek). After 24 h, 48 h, and 72 h, the MEF were infected with 10^5^ trypomastigotes per well. The slides were fixed with 10% formalin for 15 min. All samples were embedded in Vectashield with DAPI (Vector Laboratories) and covered with a 1.5 AutomatStar coverslip (DURA group). Samples were imaged using the Leica DM 5000B microscope with a Sola FISH 365 LED light source. Images were taken with a 10x ocular, plus a 20x or 63x objective, with phase contrast and DIC, respectively. Fluorescence images were taken by using the filter cubes A4 (excitation 377/50 nm, emission 447/60 nm) or L5 (excitation 470/40 nm, emission 525/50 nm).

### Macrophage isolation

Peritoneal mouse macrophages were obtained from female CD1 mice (30 – 35 g body weight) as follows: 2 ml of a 2% (wt/vol) starch solution in distilled water were injected i.p., and macropahges were harvested 24 h later by peritoneal lavage with RPMI medium containing 1% anticontamination cocktail (100 μl in 10 ml, [29]). After centrifugation at 460 g at 4 °C for 15 min, the supernatant was removed, and the pellet was resuspended in RPMI medium containing 1% anticontamination cocktail, 10% heat-inactivated fetal calf serum (iFCS) and 15% RPMI containing LADMAC (ATCC® CRL2420™) growth factors. The expanded peritoneal mouse macrophages (ePMM) were kept in this medium at 37 °C for 3-4 days and then detached with 5 min trypsin treatment and cell scrapers. The cells were counted with a Neubauer hemocytometer.

### High content microscopy

All assays were performed on an ImageXpress Micro XLS (Molecular Devices) high-content microscope. Fluorescent imaging was done using the following filter cubes: GFP (300 ms exposure), DAPI (50 ms exposure), and Cy5 (300 ms exposure). Phase-contrast images were taken on the transmitted light channel (TL10, 10% illumination). All images were taken using a 20x Zeiss objective and a cooled CCD camera with (6.45 µm x 6.45 µm pixel size, 1392 x 1040 pixel resolution).

### Live high content assay

For the live high content assay, 10^4^ ePMM were seeded into the central wells of a black 96-well plate in 100 µl of RPMI medium supplemented with 1% anticontamination cocktail [29], 10% iFCS and 15% RPMI containing LADMAC growth factors. The border wells were filled with 100 µl water. Every 24 h a new set of wells was infected with 3×10^4^/well culture-derived trypomastigotes leading to an MOI of 3 parasites to 1 host cell. This had previously been tested to lead to an optimized detection of parasites per host cells with a geometric mean of 1.1 parasites per host cells [95% confidence interval from 0.82 to 1.4]. 24 h post-infection (hpi), the remaining extracellular trypomastigotes were washed off twice with 200 µl supplemented RPMI and the infected host cells were further cultivated in 100 µl RPMI, with serial dilution of drugs. On the sixth day of infection, the plate was covered with translucent PCR film (Eppendorf AG) and placed into the ImageXpress Micro XLS microscope into an environmental chamber with 37°C, humidity, and no additional CO_2_. After 1 h acclimatization, the focus plane was determined. On 9 sites per well, images were taken every 4 h sing the GFP filter set (300 ms exposure) and transmitted light with 10% illumination (300 ms exposure).

After live imaging, all supernatant was removed and the cells were fixed with 10% formalin for 15 min at room temperature and stained with 100 µg/ml Hoechst 33342 (Merck) for 30 min in the dark at room temperature. The plate was stored at 4 °C until it was imaged with the ImageXpress using transmitted light with 10% illumination (300 ms exposure), the GFP (300 ms exposure) and the DAPI (50 ms exposure) filter set on 9 sites per well.

### Image analysis

Image analyses were performed on the MetaXpress 6 software. For live imaging, green fluorescent parasites were quantified from the GFP channel. The TopHat filter was applied with 10 pixel diameter. Round objects of size 1-10 µm and a fluorescence difference of over 1000 were designated as parasites (from the filtered image). For imaging of fixed cells, host nuclei and parasite kinetoplasts were counted. Parasites’ mitochondrial DNA (kinetoplast DNA, kDNA) is AT-rich and was therefore stained more strongly using minor-groove binding stains such as Hoechst 33342 [30]. Round objects of size 5 – 30 µm, with a fluorescence difference of over 1000 were designated as host cell nuclei. Parasite kDNA was detected by applying a TopHat filter with 10 pixels diameter as round objects of 1-10 µm and a fluorescence difference of over 1000, which did not coincide with host cell nuclei. Parasite kDNA in an area where the green fluorescence difference was lower than 1000 were defined as GFP-negative parasites. The remaining parasites were denoted GFP-positive parasites.

### Statistical analyses

Statistical analyses were performed in R version 3.5.1 [31] using the packages “tidyverse” [32], “readxl” [33], and “viridis” [34]. Scripts for different analysis steps are available under GitHub (https://github.com/fesser-af/NiMDA). The dose-response relationship for each time point of drug exposure was quantified by a four-parameter log-logistic model (Equation 1) using the package “drc” [35] in R.

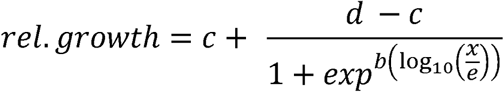

*Equation 1 Dose-response model. The four-parameters of this model were the upper plateau (d), the lower plateau (c), the hill slope (b), and the inflection point (e). The inflection point (e) corresponds to the half maximal effective concentration (EC50)*.

Exponential models of change in parasite numbers (Equation 2) were determined for each site in R.

Exponential multiplication was assumed to dominate the replication period.

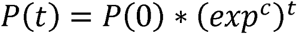

*Equation 2 Exponential model for change in parasite numbers. P(t)is the number of detected parasites at a certain time after infection (in h), P(0) is the hypothetical number of parasites at this site assuming an exponential growth over the whole range of the time after infection, exp(c) describes the fold change in parasite numbers per h, t is the time of infection of interest (in h)*

## Results

### Transfected *T. cruzi* expressing eGFP in the replicating life-cycle stages

To enable fluorescence-based live imaging, *T. cruzi* STIB980 epimastigotes were transfected with a linearized plasmid containing the *eGFP* gene (enhanced green fluorescent protein) under control of the rRNA promoter (Fig 1 A), which was designed to integrate into the spacer region within the genomic rRNA locus. Fluorescent metacyclic, amastigote and trypomastigote forms were derived from the transfected and cloned epimastigote transformants as described (Methods). eGFP expression was detected in the replicating stages of the transfected parasite line by flow cytometry and epifluorescence microscopy (Fig 1 B-F). As quantified by flow cytometry, the green fluorescence levels (excitation 488 nm, emission 522 nm) in epimastigote forms were about 100 times higher than the autofluorescence levels of non-transfected cells (Fig 1 B). Epifluorescence imaging showed an even distribution of eGFP throughout the cytosol in epimastigotes and amastigotes, the two replicating stages (Fig 1 C and E, respectively). Green fluorescence could also be detected in the flagellum of epimastigotes and the short flagellum of amastigotes. In contrast, in the non-replicating stages, the metacyclic trypomastigotes and trypomastigotes (Fig 1 D and F, respectively), green fluorescence was barely distinguishable from the autofluorescence of the non-transfected parent.

When cultivated for 3 months in the absence of antibiotic selection, only about 50% of the transgenic *T. cruzi* amastigotes still expressed eGFP at a detectable level (data not shown). When epimastigote forms of the transgenic *T. cruzi* line were cultivated without antibiotics for six months, about 95% still expressed eGFP (data not shown). For this reason, amastigote cultures were only maintained for 4 weeks after they had been derived from epimastigotes.

### A new assay design for live monitoring of drug action

We have devised a new plate design combined with a special imaging scheme that enables observation of parasite development over 6 days, with only 24 h of user time on the high-content microscope (Fig 2). The mammalian cells were cultured in 96-well plates, but were not all infected with *T. cruzi* at the same time (Fig 2 B). Instead, every 24 h, a new row of 10 wells was infected with an MOI of 3:1 (Fig 2). 24 h post infection (hpi), the wells were washed thoroughly to remove extracellular parasites. Then, 7 wells were treated with a 3-fold serial dilution of test compound, while the remaining wells were left untreated. Thus in every row on the plate, the parasites had the same period of infection and the same period of drug exposure. In every column, the parasites were exposed to the same drug concentration. By day 6, when all the rows were used up, the infection period covered 1 to 6 days and drug treatment from 0 to 5 days. The plate was then placed into the microscope for automated live imaging over 24 h. Every 4 h, images were taken from 9 sites per well on the GFP channel for parasite quantification as well as with transmitted light for quality control. After 24 h, the plate was fixed with 10% formalin and stained with Hoechst 33342. All 9 sites per well were imaged once again on the DAPI channel, GFP channel, and with transmitted light. The DNA stain enabled the number of host cells to be determined by nuclei count (5 – 30 µm round objects), with the number of parasites inferred from counting their kinetoplasts (1 – 10 µm round objects). The parasite kinetoplast is brighter than the parasite nucleus, because the AT-rich kinetoplast DNA binds Hoechst preferentially [30].

**Fig 2.**
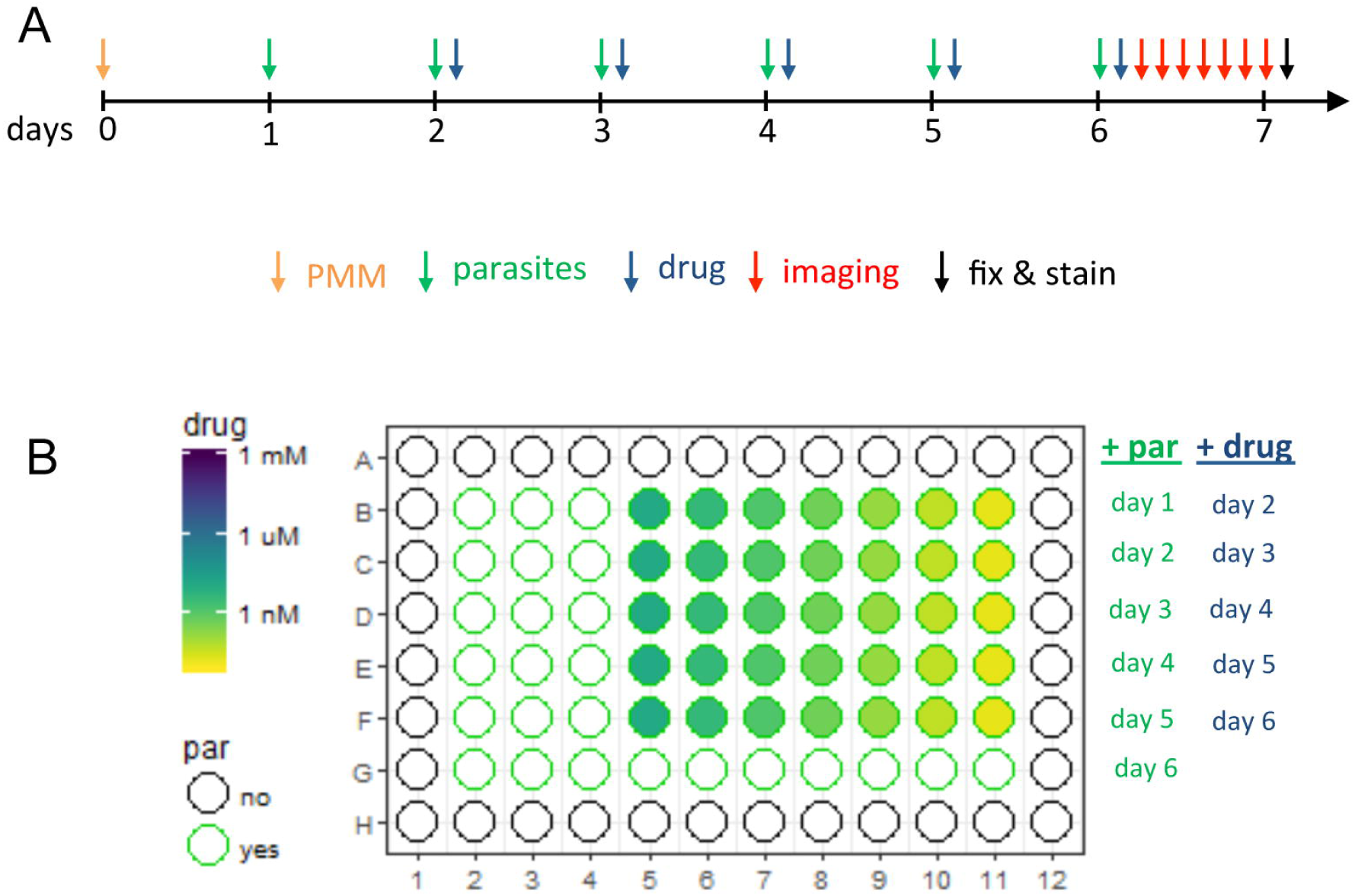
Assay design. The timeline of the experimental set-up (A) and plate design (B). On a 96 well plate, a new set of wells containing expanded peritoneal mouse macrophages (ePMM) was infected every 24 h. 24 h hpi, extracellular parasites were removed and drugs were added in 3-fold serial dilution. On day 6, after addition of parasites and drugs (either benznidazole or posaconazole), the plate was imaged on 9 sites per well over 24 h every 4 h (live imaging). After 24 h live imaging, the plate was fixed and stained with Hoechst 33342 and imaged again (fixed imaging). Green fluorescent parasites per image were detected for all images (from live and fixed imaging) of the green fluorescent channel. Kinetoplasts per image were detected on images of the DAPI channel (fixed imaging).

### High content imaging reveals the high degree of variability in untreated cultures

All assays were performed as biological triplicates, which resulted in a total of 27,540 fluorescent microscopy images per drug tested. On every image of the GFP channel, the number of green fluorescent parasites was determined; on the DAPI channel images of fixed cells, the numbers of host nuclei and parasites (on basis of kDNA) were also determined.

Parasite numbers per image were driven by the time post infection, drug concentration and the drug exposure time (Fig 3). There was a high degree of variability: between the days of infection (rows on the plate), between replicate wells in the same row, and even between different sites in the same well. Nevertheless, there were common trends in the dynamics of infection (Fig 3 A and B). In the untreated control cultures, the increase in parasite numbers over the period of infection (6 days) could be separated into three phases. During the first 24 h, the increase was usually the most pronounced, but also displayed the highest variance, consistent with settling of parasites into the focal plane and the differention of trypomastigotes that barely expressed eGFP to amastigotes that express eGFP. During days 2 to 4, parasite numbers, as determined by green fluorescence, continued to increase, consistent with amastigote replication, but began to level off towards day 4 (Fig 3 A). On days 5 and 6, parasite numbers inferred from the green fluorescent channel increased only slightly, if at all (Fig 3 A). Around day 4, differentiation to trypomastigotes dominates the development in parasite numbers over amastigote replication. Fixation of cells allowed a direct comparison between the parasite numbers determined by green fluorescence and those determined from DNA-staining of the kinetoplast (Fig 3 A and B and Supp Fig 1). The latter method returned higher numbers. Parasites detected as kDNA on the DAPI channel, but not on the GFP channel, could be either trypomastigotes that did not express eGFP, dead parasites whose kinetoplast was still intact, or revertants that no longer express the *eGFP* gene. During the middle phase (day 3 post infection), the proportion of parasites detected on the green fluorescent channel to parasites detected on the DAPI channel was 0.95. This suggests that the plateauing of GFP-positive parasites at later time points was due to the transformation of intracellular amastigotes to trypomastigotes and not to a loss of the *eGFP* gene.

**Fig 3.**
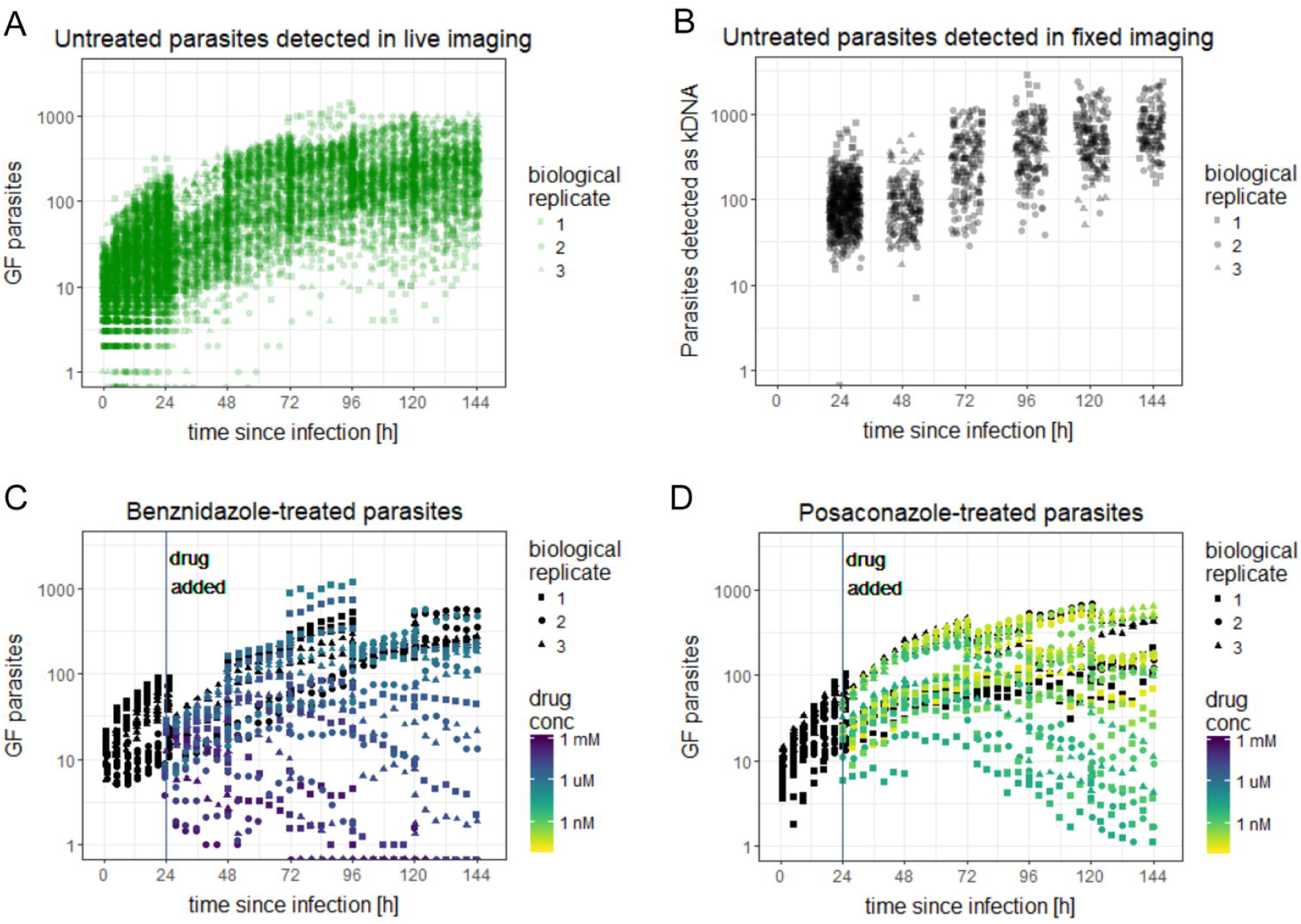
Parasite numbers over time after infection and drug exposure. The assay design was used to monitor the development of parasite numbers for posaconazole and benznidazole treated parasites. Number of untreated parasites per image over time after infection are depicted from live (B) and fixed (C) imaging with detection as green fluorescent parasites or as kinetoplasts, respectively. Mean number of treated parasites per image per well is depicted for benznidazole (D) and posaconazole (E) in live imaging detected as green fluorescent parasites. Untreated controls are depicted in black (D, E).

### The decrease in EC50 values over time of drug exposure is a characteristic feature of a drug

For each time point of drug exposure, the growth relative to the untreated control at the same stage of infection depends on the drug concentration. A four-parameter log-logistic model was used to determine this relationship and the half maximal effective concentration (EC50, Fig 4 A and B).

**Fig 4.**
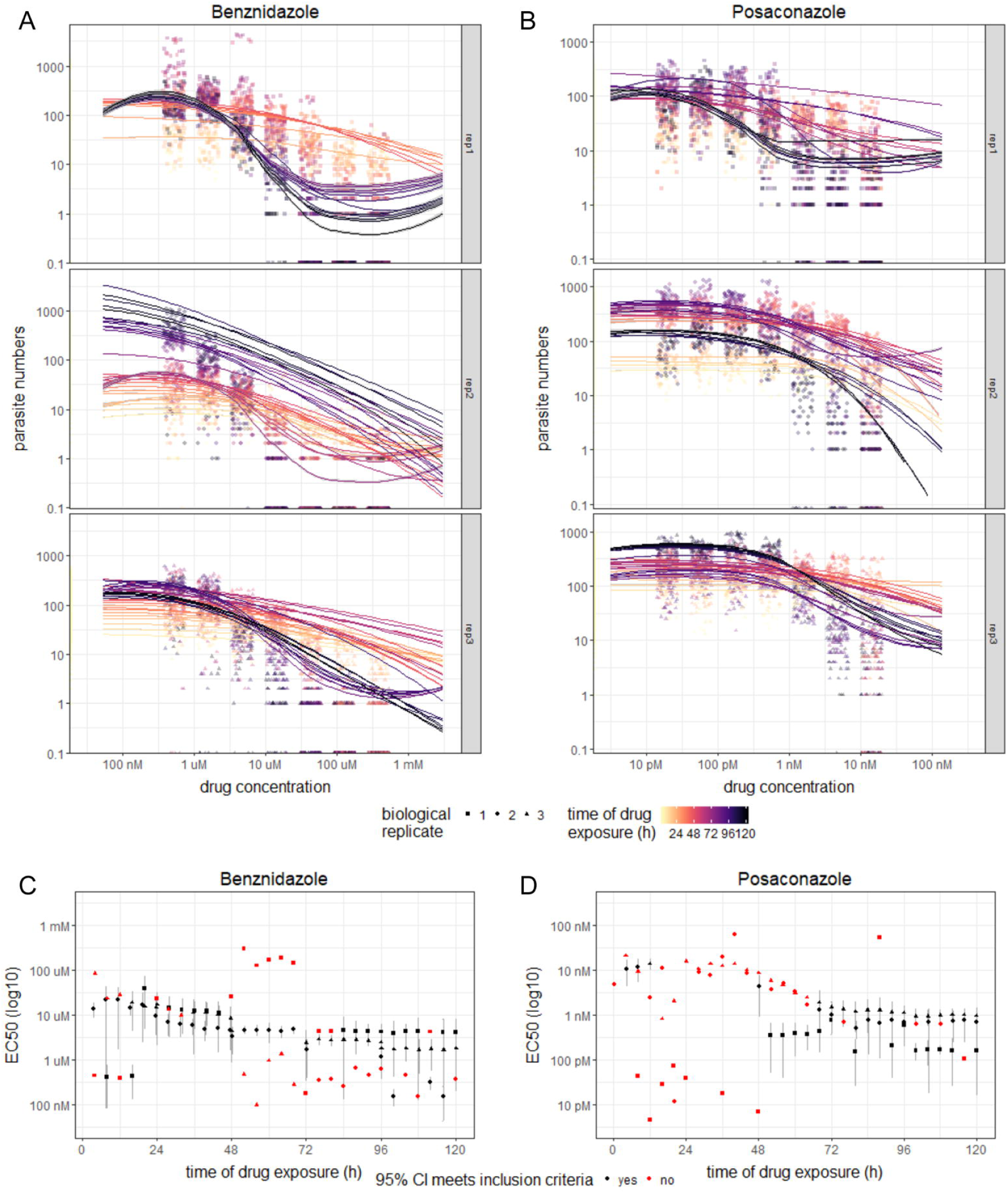
Development of the concentration of half-maximal effect (EC50) over time of drug exposure. Dose-response curves were estimated for each time point after drug exposure by modelling growth relative to the same-aged untreated control using the “drc” package in R for benznidazole-(A) and posaconazole-treated (B) parasites. The resulting estimates of the EC50 are plotted over time of drug exposure (C, D). For several time points the 95% confidence interval of the EC50 value did not meet the inclusion criteria: to be between 30 nM and 3 mM for benznidazole and to be between 300 nM and 3 pM for posaconazole.

Over the time of drug exposure, the EC50 values decreased until they reached a stable low point (Fig 4 C and D). How soon the final EC50 value was reached, was as characteristic for a drug as the final EC50 value itself. For benznidazole, the EC50 value decreased from a median value of 17 µM (with 95% confidence intervals ranging from 2.4 µM to 70 µM) after 24 h of drug exposure to 3.8 µM (with 95% confidence intervals ranging from 0.87 µM to 7.9 µM) after 48 h of drug exposure. Afterwards, the majority of the EC50 values remained in this range. In contrast, for posaconazole, during the first 48 h, only 4 EC50 values could reliably be determined with a median of 11 nM (95% confidence intervals ranging from 0.88 nM to 18 nM). 68 h after drug exposure, an EC50 value could be determined in all replicates for the first time with a median value of 1.3 nM (95% confidence intervals ranging from 0.15 nM to 3.4 nM). Afterwards, the majority of the EC50 values remained in this range.

The EC50 values in these experiments were highly variable, even between biological replicates. At some time points, the EC50 values could not be determined with a reasonable 95% confidence interval. This is consistent with the fact that the parasite numbers have a high variability even in the untreated culture. Thus, classical end-point read-outs such as EC50 may not be the most suitable method for quantifying drug action with live imaging. We therefore aimed for a more robust read-out.

### The change in parasite numbers at each site can be quantified in an exponential model

At every imaged site, the same set of host cells was observed over the 24 h period of imaging. Therefore, for each site, the change in parasite numbers between time points in this 24 h interval (as exemplified in Fig 5 A) only depended on the age of infection, the drug concentration and the time of drug exposure. We determined an exponential model for parasite numbers over time at every imaged site (Methods, Equation 2).

**Fig 5.**
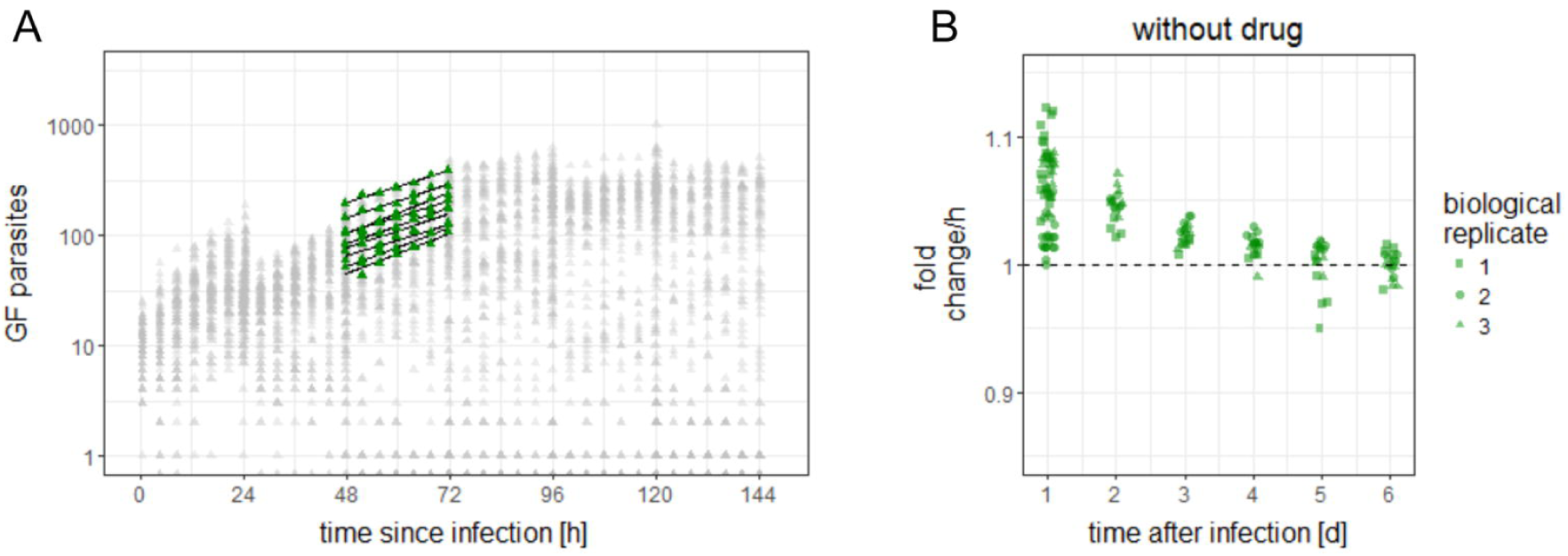
Exponential model of the development of parasite numbers over time. Numbers of green fluorescent parasites from one plate are plotted over time after infection (A). The parasite numbers obtained at the seven time points from the 9 sites in one well from this plate are highlighted in green as an example. The exponential models (Methods) determined for every site of this well are plotted for illustration. The variable “fold change per h” of those exponential models for every site of all plates changes between days after infection (B).

This exponential model of change in parasite numbers has two parameters: the offset P(0) and the slope c (Equation 2). In the logarithmized version of the equation, the slope c can be negative, zero or positive, corresponding to declining, stagnating and growing parasite numbers, respectively. The slope c is a net result of replication rate, differentiation rate and parasite reduction rate. The offset (basal parasite number, P(0)) is an extrapolation of the number of parasites in the monitored region at the time of infection.

Most of the observed variability in parasite numbers was captured by the variability in P(0), which already differed between different sites of one well. This was to be expected since the numbers of infected host cells are not evenly distributed over the whole well, and some infected host cells would have been infected by more than one parasite.

While the extrapolated basal parasite number P(0) was very variable, the fold change in parasite numbers (i.e. the exponential of the slope, exp(c)) was more robust, especially between days 2 – 4. In addition, the fold change is more informative as it directly reflects the difference between replicating and dying amastigotes (at least in the early phase, when the amastigotes do not differentiate to trypomastigotes; Fig 5 B). For these reasons, we used the fold change of parasite numbers over time of drug exposure to characterize drug action. In particular, we focused on the tipping point of drug action, i.e. the time point when the parasites had stopped growing and started to die.

### The tipping point of drug action is a sensitive and robust readout

The fold change in parasite numbers in drug-treated cultures was calculated based on the exponential model of parasite replication. Initially, the parasites continued to replicate, although often more slowly than in the untreated control. After some time of drug exposure, at certain concentrations, parasite numbers start to decrease, i.e. parasites are dying. Once most parasites are killed, the parasite numbers will not decrease further and the fold change will approximate to 1. If the fold change exceeds 1 again, this might indicate that surviving parasites are replicating, or reflect an issue with drug stability. Certain drugs might be cytostatic at some concentrations. In this case, the net fold change would be stable at approx. 1 over the whole period of drug exposure.

The tipping point was defined as the time point immediately after the net fold change dropped below 1. The tipping point of drug action is both time- and concentration-dependent. Fig 6 shows the fold change in parasite numbers for each concentration and for every day post drug exposure. For each replicate and each drug concentration, the tipping point, i.e. the day when the fold change in parasite number has significantly dropped below 1 (95 % confidence interval excluding 1), is marked.

**Fig 6.**
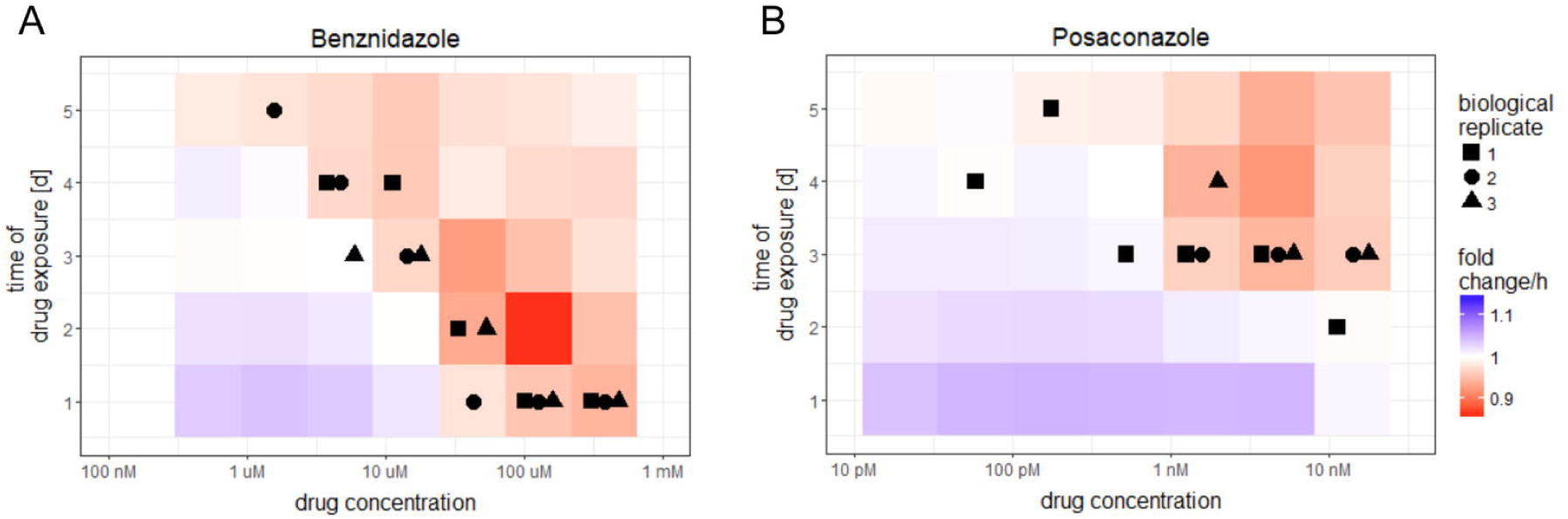
The fold change in parasite numbers of drug-treated parasites depends on drug concentration and time of drug exposure. The fold change in parasite numbers was obtained by exponential models of change in parasite numbers for every site (Methods). The fold change in parasite numbers depended on the concentration and time of exposure with benznidazole (A) and posaconazole (B). The first day of drug exposure, at which the fold change was significantly below 1, is indicated for every replicate.

In the benznidazole-treated wells, parasite numbers started to decrease within the first 24 h of drug treatment at the highest concentrations. After 48 h, parasite numbers reached 0 for the highest concentration. In the posaconazole-treated wells, parasite numbers continued to increase for at least 2 days. For benznidazole, there was a correlation between drug concentration and the tipping point of drug action (Fig 6 A). In contrast, in most cases, the posaconazole-treated parasites only started dying during the third day of drug exposure, even at the highest concentration (Fig 6 B). In summary, calculating the tipping points allowed the measurement of time- and concentration-dependence of drug action against intracellular *T. cruzi* amastigotes, and illustrated potential shortcomings of posaconazole as compared to benznidazole.

## Discussion

High-content microscopy has led to great innovation in kinetoplastid drug discovery [6, 9-14, 36]. It combines the high throughput capacity of systems such as the LacZ assay [12, 37], with the more detailed information than is obtainable by manual counting [15]. Nuclear staining as a read-out allows high-throughput *in vitro* assays to be undertaken with a broad panel of strains [6]. However, the use of nuclear staining generally requires fixation of the cells, precluding live imaging over several days to observe the time-course of drug action. Live imaging can be more easily done with fluorescent *T. cruzi* reporter lines.

We established an eGFP expressing parasite line for use in live imaging assays. This parasite line expresses eGPF in high levels from the ribosomal locus, but only in the replicating life-cycle stages. This is in line with the observation that the reduction of transcription in the non-replicating (metacyclic) trypomastigote stage is particularly pronounced for ribosomal loci [38].

The new assay design presented here enables the monitoring of *T. cruzi* amastigote replication for 6 days post-infection, and drug action over 5 days of exposure, in four-hourly intervals. Nine sites per well can be monitored separately. This not only creates a wealth of data – it also reveals a high degree of variability in *T. cruzi* numbers, not only between different wells and different days of infection, but also between different sites in the same well, infected on the same day. This complicated the EC50 determination at each time point.

In contrast, the fold change per h in the number of parasites over the period of 24 h after drug incubation is a sensitive and robust measure of pharmacodynamics. The fold change on a specific day of infection was very repeatable within wells, between wells, and between days of infection. The fold change is influenced by the drug concentration and the time of drug exposure. For each drug concentration, we can determine the day of drug exposure at which the net fold change per h drops significantly below one. At this tipping point, more parasites are dying than replicating, providing a statistically robust quantitative measure of the time-to-kill. The graphical representation of the fold change per h of parasite numbers in relation to drug concentration and exposure time clearly depicts the different pharmacodynamics of benznidazole and posaconazole. For benznidazole, the time-to-kill decreases roughly exponentially with increasing drug concentration. In contrast, even at the highest concentrations of posaconazole tested, the net killing only starts after 3 days of drug exposure. We propose to introduce the concentration-dependency of time-to-kill as a novel measure to benchmark drug candidates for Chagas’ disease.

## Acknowledgement

Isolation and expansion of peritoneal macrophages was performed by Romina Rocchetti. Monica Cal provided cell and parasite culture support and support with high-content microscopy. Dr. Remo Schmidt supported the molecular biology and basic statistic analyses.

We kindly thank Prof. Dr. Till Voss and his group for letting us use the high-content microscope.

Special thanks to him and Dr. Nicolas Brancucci for fruitful discussions on high-content use.

We thank Prof. Dr. Antonio Osuna for the kind provision of the *T. cruzi* strain.

We thank Prof. Dr. Joachim Clos for the kind provision with the LADMAC cells.

This work would not have been possible without the financial support of the Swiss National Science Foundation, the Nikolaus und Bertha Burckhardt-Bürgin-Stiftung, and the Freiwillige Akademische Gesellschaft.

Olivier Braissant’s work is supported by the Merian Iselin Stiftung

## Supporting informations captions

Supp. Fig 1. **Development of parasite numbers over time for all replicates separately**. Parasite numbers per image from untreated wells of all biological replicates from live imaging (green, detected as GFP positive parasites) and fixed imaging (black, detected as kinetoplasts).

## Author contributions

Conceptualization: AFF, MK, PM

Formal Analysis: AFF, OB

Funding Acquisition: MK, PM, AFF

Methodology: AFF, OB, FO, JMK, MK

Resources: FO, JMK

Supervision: MK, PM

Visualization: AFF

Writing – Original Draft Preparation: AFF, PM

Writing – Review & Editing: AFF, OB, FO, JMK, MK, PM

## Notes

https://github.com/fesser-af/NiMDA

