## Supplementary figures and images for "Non-invasive monitoring of drug action: a new live *in vitro* assay design for Chagas’ disease drug discovery"

### Supplemtary figure 1

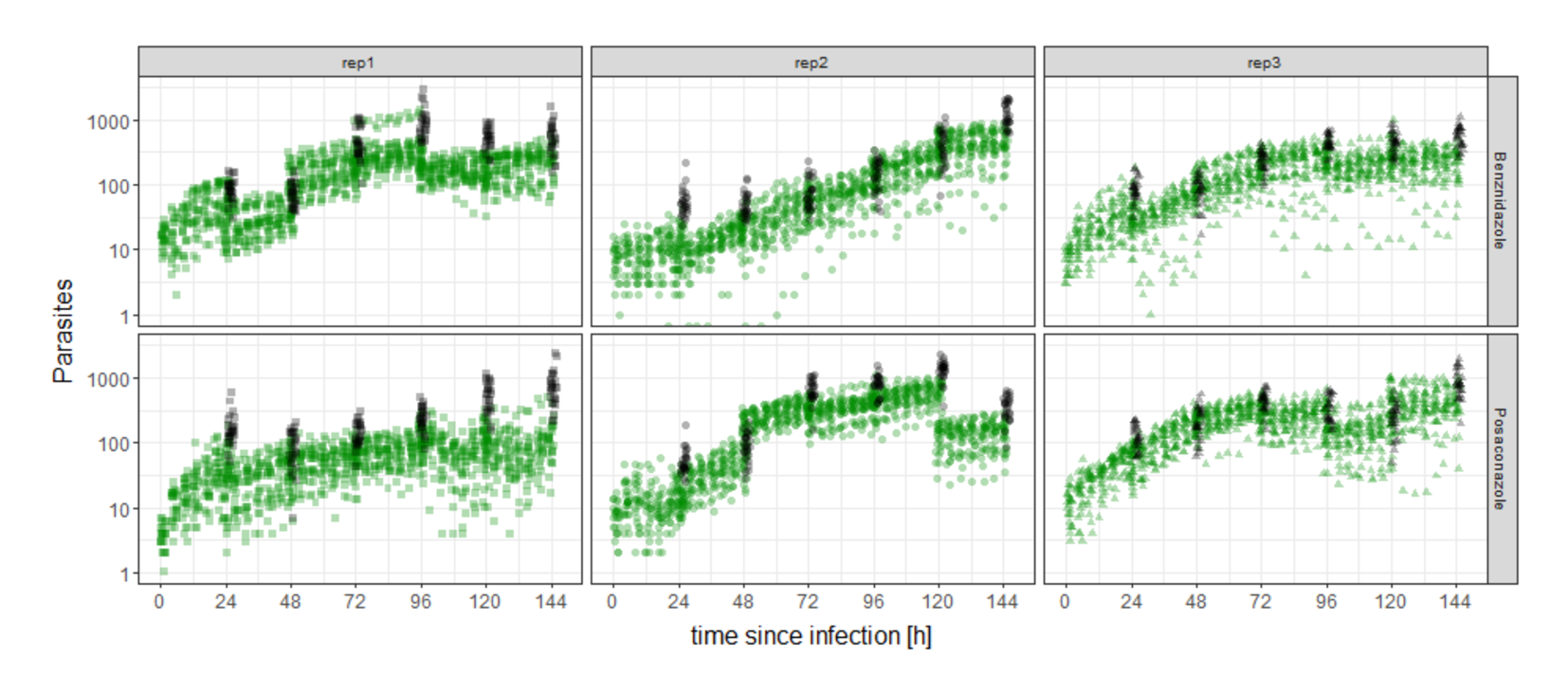
